# Integrative analysis of single cell genomics data by coupled nonnegative matrix factorizations

**DOI:** 10.1101/312348

**Authors:** Zhana Duren, Xi Chen, Mahdi Zamanighomi, Wanwen Zeng, Ansuman T Satpathy, Howard Y. Chang, Yong Wang, Wing Hung Wong

**Author notes:** Authors contributed equally to this work (Zhana Duren, Xi Chen, Mahdi Zamanighomi). Corresponding author: Wing Hung Wong, Mailing Address: Department of Statistics, Sequoia Hall, 390 Serra Mall, Stanford University, Stanford, CA 94305-4065.

## Abstract

When different types of functional genomics data are generated on single cells from different samples of cells from the same heterogeneous population, the clustering of cells in the different samples should be coupled. We formulate this “coupled clustering” problem as an optimization problem, and propose the method of coupled nonnegative matrix factorizations (coupled NMF) for its solution. The method is illustrated by the integrative analysis of single cell RNA-seq and single cell ATAC-seq data.

**Significance Statements:** Biological samples are often heterogeneous mixtures of different types of cells. Suppose we have two single cell data sets, each providing information on a different cellular feature and generated on a different sample from this mixture. Then, the clustering of cells in the two samples should be coupled as both clusterings are reflecting the underlying cell types in the same mixture. This “coupled clustering” problem is a new problem not covered by existing clustering methods. In this paper we develop an approach for its solution based the coupling of two nonnegative matrix factorizations. The method should be useful for integrative single cell genomics analysis tasks such as the joint analysis of single cell RNA-seq and single cell ATAC-seq data.

## Introduction

Biological samples of interest in clinical or experimental studies are often heterogeneous mixtures, i.e. a sample may consist of many different subpopulations of cells with distinct cellular states. To resolve the heterogeneity and to characterize the constituent subpopulations, it is necessary to generate functional genomic data at the single cell level. An exciting recent development in genomics technology has been the emergence of methods for single cell (sc) measurements, for example, scRNA-seq (1) enables transcription profiling, scATAC-seq (2) offers chromatin accessibility data, sc-bisulfite sequencing (3) measures DNA methylation, all at the single cell level.

Often, the first step in the analysis of single cell data is clustering, which aims to resolve the single cells into the constituent subpopulations. Clustering methods for scRNA-seq data were discussed in (4, 5), and clustering of scATAC-seq data were given in (6). Existing methods, however, do not cover the increasing common situation when two or more types of sc-genomics experiments were performed on different subsamples from the same cell population. For example, Figure 1A depicts the situation when one subsample is analyzed by scRNA-seq while another is analyzed by scATAC-seq. Since both types of measurements are informative about the constituent cell types in the heterogeneous population, it is clear that we should aim to couple the two clustering processes in such a way that the clustering of the cells in the scRNA-seq sample can also make use of information from the scATAC-seq sample, and vice versa. In this paper we formulate this “coupled clustering” problem as an optimization problem, and introduce a method, called coupled nonnegative matrix factorizations (coupled NMF), for its solution.

**Fig. 1.**
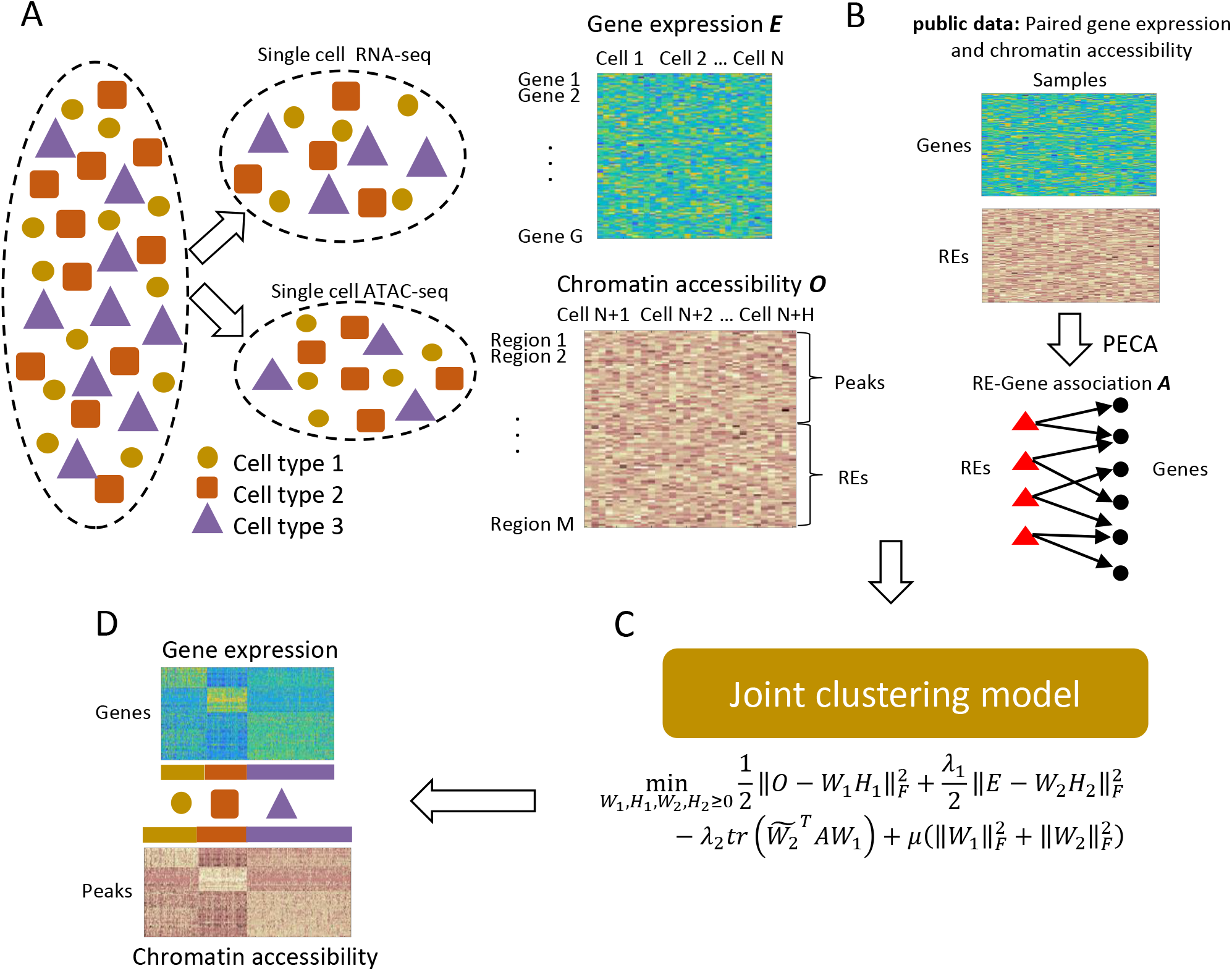
Overview of the coupled clustering method.

We first introduce our approach in general terms. Let *O*be a *p_1_* by *n_1_* matrix representing data on *p_1_* features for *n_1_* units in the first sample, then a “soft” clustering of the units in this sample can be obtained from a factorization *O=W_1_H_1_* as follows: the *i^th^* column of *W_1_* gives the mean vector for the *i^th^* cluster of units, while the *j^th^* column of *H_1_* gives the assignment weights of the *j^th^* unit to the different clusters. Similarly, clustering of the second sample can be obtained from the factorization *E=W_2_H_2_* where E is the *p_2_* by *n_2_* matrix of data on *p_2_* features (which are different from the features measured in the first sample) for *n_2_* units. To couple two matrix factorizations, we solve the optimization in Figure 1 C, where the cost function contains a term 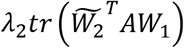 where 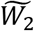 is a submatrix containing a subset of rows of *W_2_*, and *A* is a “coupling matrix”. The construction of *A* is application specific but depends on the assumption that, based on scientific understanding or prior data, it is possible to identify a subset of features in one of the sample that are linearly predictable from the features measured in the other sample. In such a situation, we can take *A* to be the matrix representation of the linear prediction operator. For the application of interest in the current paper, where gene expression is measured in one sample and chromatin accessibility is measured in the other sample, we will use a diverse panel of cell lines with both expression and accessibility data, to train a prediction model of gene expression from accessibility. See Approach section for details.

## Approach

### Construction of Data Matrices

From single cell ATAC-seq data, we compute a data matrix 0, where *O_ij_* denotes the degree of openness (i.e. accessibility) of the *i^th^* region in the *j^th^* cell (6). By region we mean union of predefined regulatory elements (REs) and peaks. From single cell RNA-seq data, we compute the data matrix *E*where *E_gh_* denotes the expression level of the *g^th^* gene in the *h^th^* cell (7). Details are given under data processing in the Methods and Materials. Note that the single cell ATAC-seq and the single cell RNA-seq data are not measured in the same cell (Fig. 1 A).

### Construction of coupling matrix

Our approach to the initialization of *A* is to look for a subset of genes whose expression is highly predictable from chromatin accessibility of nearby REs. To do this, we take advantage our recent work on modeling paired gene expression and chromatin accessibility data (on bulk samples) across diverse cellular contexts (8). From the PECA model in that work, for each gene g, we can extract a set *S_g_* of REs that regulate that gene. We consider the regression model of target gene (TG) expression (denoted as *E_g_*) on its regulatory elements’ (RE) accessibility (denoted as *O_i_*).

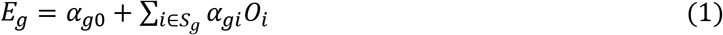

We estimate the parameter α_g_ by fitting the penalized least square problem (*eq*. 2 below) based on expression and accessibility data on a diverse panel of cell lines (56 cell lines in the case of mouse, and 148 cell lines in the case of human (Supplementary Table 1).

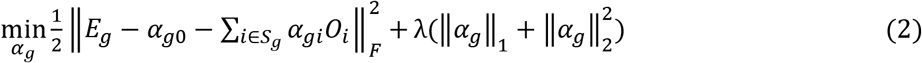

where λ is determined by 5-fold cross validation. After fitting the model, we select a set of “well predicted” genes (denoted as *S*) for which the relative prediction error is less than 0.3, that is, 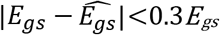, for at least 80% of the cell lines in the panel. In this way, we selected 5,281 well predicted genes in mouse and select 6,537 well predicted genes in human. The initial value of the coupling matrix *A* is obtained from the coefficients α_g_ associated with these well predicted genes.

### Clustering by Coupled nonnegative matrix factorizations

The gene set *S*and the matrix *A* allow us to couple expression-based clustering and accessibility-based clustering according to a regulatory model supported by extensive prior data. Once the coupling is defined this way, we can obtain the factorizations of the two data matrices by solving the following optimization problem (Fig. 1*C*).

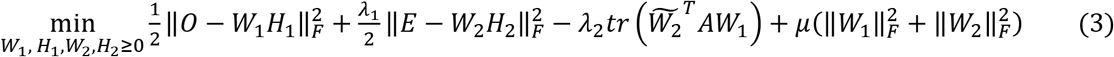

Where 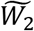 represent a submatrix of *W*_2_ only containing the rows corresponding to the genes in *S*, the first row of 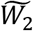 is [1,1,…1], which corresponding to the first column of matrix *A* [α_10_, α_20_, … α_n0_]^T^, the intercept of the regression model in *eq. (1)*. After solving the optimization, the cluster profile and the cluster assignments for the *k^th^* cluster in the accessibility data can be obtained respectively from the *k^th^*column of *W_1_* and the *k^th^* row of *H_1_*. Similarly, the expression-based clustering can be obtained from *W_2_* and *H_2_* (Fig. 1*D*). In this application *AW_1_* gives the cluster-specific predictions of the expression of genes in *S*based on the cluster-specific accessibilities of REs, and hence the trace term enforces our expectation that the expression of genes in S should be consistent with the predictions based on accessibility of nearby REs. We can refine the coupling iteratively, as follows. We assign single cells to clusters according to the assignment weights given by *H_1_* and *H_2_*. After getting the cluster results, we choose cluster-specific genes based on single cell RNA-seq clustering. Then we restrict the gene set *S* in cluster-specific gene and re-cluster the cells by optimizing the objective function in *eq*. (3). We continue until the cluster assignments are not changed by further iterations.

## Results

### Results on simulation data

The performance of our coupled clustering method is first evaluated in a simulation study where single cell data is simulated by mixing reads that are sampled from two bulk data sets corresponding to two cell types. The bulk data sets used in our simulation study are from two very similar cell types from hematopoietic differentiation process, namely common myeloid progenitor (CMP) and megakaryocyte erythroid progenitor (MEP) (9). We randomly sample 3,000 reads from bulk RNA-seq data and add a Gaussian noise of SNR = 5 to simulate a single cell RNA-seq. Similarly, we randomly select 40,000 reads from bulk ATAC-seq data and add a Gaussian noise of SNR = 5 to simulate a scATAC-seq data. Number of reads in our simulation data are similar to the number of reads in 10 X genomics scRNA-seq and C1 Fluidigm scATAC-seq data. In our simulation, we generated 100 scRNA-seq and 100 scATAC-seq for CMP and MEP respectively.

To simulate a scRNA-seq data set from a mixed population with two cell types, we simply combine the 200 single cell RNA-seq data from two cell lines together and treat it as a single scRNA-seq data set. We then apply k-means and non-negative matrix factorization (NMF) to cluster the mixed cells. We run k-means 50 times with different random initial values and choose the result that gives the minimum total sum of within cluster distances. Similarly, we run NMF 50 times and choose the result that gives the minimum approximation error in Frobenius norm. The results of all the 50 runs on scRNA-seq and scATAC-seq data by k-means and NMF are shown in Supplementary Fig. S1. Finally, we perform coupled NMF clustering based on both the scRNA-seq sample (200 cells mixture) and the scATAC-seq sample (200 cells mixture). The performance of the three clustering results (k-means on scRNA-seq only, NMF on scRNA-seq only, and coupled NMF on both scRNA-seq and scATAC-seq) are presented in Figure 2A). A similar improvement is also seen in the clustering results on the scATAC-seq sample (Fig 2B). It is seen that coupling leads to greatly improved results, reducing the assignment error rate by more than 4 folds over the other two methods (Fig 2C).

**Fig. 2.**
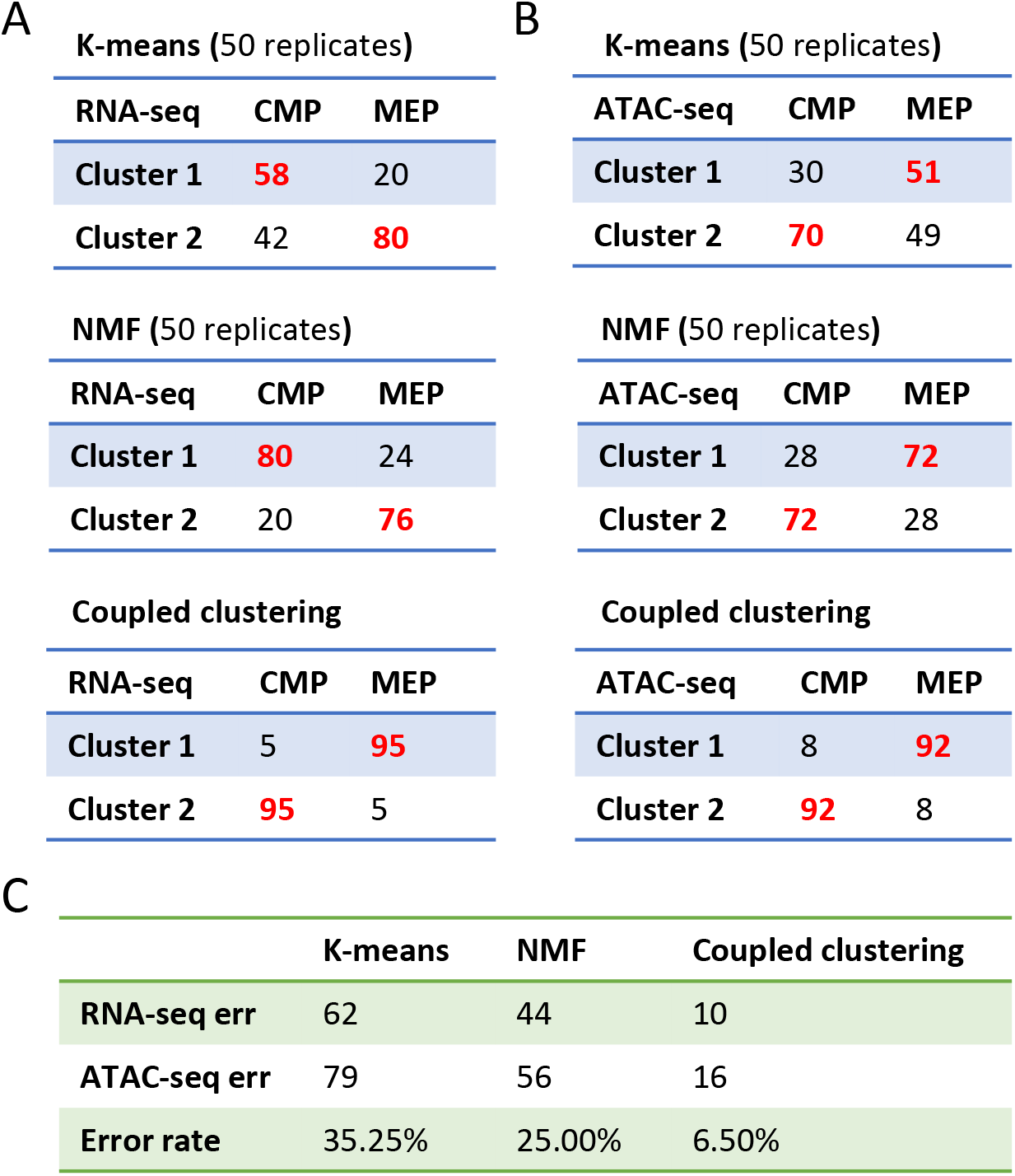
(A) Clustering results of k-means, NMF, and our coupled clustering on simulation scRNA-seq data on CMP and MEP. (B) Clustering results of k-means, NMF, and our coupled clustering on simulation scATAC-seq data on CMP and MEP. (C) Comparison of k-means, NMF, and coupled clustering on simulation data of CMP and MEP.

### Assessment of prediction model before coupling

We are interested in applying coupled NMF to analyze data generated from differentiation of mouse embryonic stem cell, namely scRNA-seq and scATAC-seq at day 4 after retinoic acid (RA) treatment (Methods and Materials). To assess whether the model learned from the diverse panel still have good predictive power in this new biological context, we first generated bulk RNA-seq and ATAC-seq from this context (i.e. RA day-4). Using the model trained on the diverse panel, we predicted the expression at RA day-4 of genes in gene set *S* based on the accessibility data at RA day-4. Figure 3 presents the observed versus predicted scatter plot, which shows the genes in S can indeed be predicted with high accuracy in this context (R^2^=0.86, r=0.93). This gives us confidence in using the model to initiate the coupling.

**Fig. 3.**
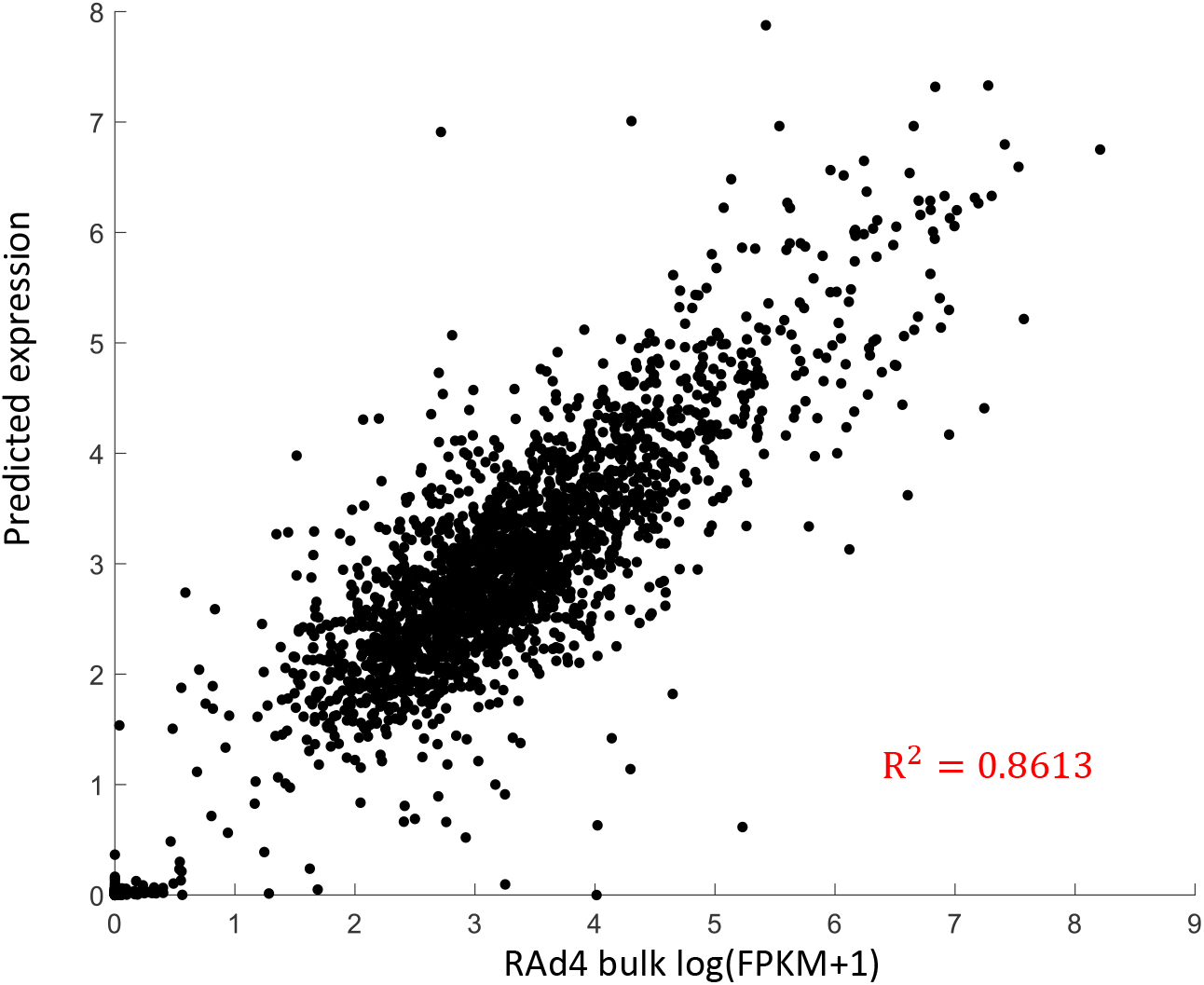
Scatter plot of RA day 4 gene expression (log(FPKM+1)) verses predicted expression values from our model.

### Results on real single cell data

Next, we test our method on scRNA-seq and scATAC-seq data generated from the RA day-4 cell population. We first perform coupled NMF with K = 2 (i.e. 2 clusters) and then visualize the clustering result on a t-SNE plot computed based on Spearman correlation. There are clearly 2 separated clusters in both the t-SNE plot from scATAC-seq and the t-SNE plot from scRNA-seq (Supplementary Fig. S2). When we increase the number of clusters to 3, we can see 3 well separated clusters in t-SNE plot. However, when K is increased to 4 or 5, the resulting clusters are no longer clearly separated (Supplementary Fig. S2). Thus we conclude that K=3 is sufficient for this data.

For each of the 3 clusters, we identify cluster-specific TFs based on their expression from RNA-seq data, and compare their motif activities across different clusters (Figure 4). Here the motif activity of a TF reflects the degree of enrichment of the TF’s motif on accessible REs (see Methods and Materials). Figure 4B shows the motif activities and expressions of some cluster-specific TFs on the t-SNE visualization (Figure 4B, Supplementary Fig. S4). Fig. 4 C shows the heat maps of motif activities and expressions for a subset cluster specific TFs, namely those with expression TPM greater than 10 in at least 40 cells. It is seen that cluster-1-specific TFs’ (e. g. Ebf1, Lhx1, and Neurod1) have high motif activities in cluster-1 specific peaks. Similarly, cluster-2-specific TFs, Gata4, Foxa2, and Jun, have high motif activity in cluster 2 and cluster-3-specific TFs, Rfx4, Sox2, Sox9, Pou3f2 & Pou3f4, have high motif activity in cluster 3. This result shows that our method leads to highly consistent TF expression and TF motif activities within each of the inferred constituent subpopulations.

**Fig. 4.**
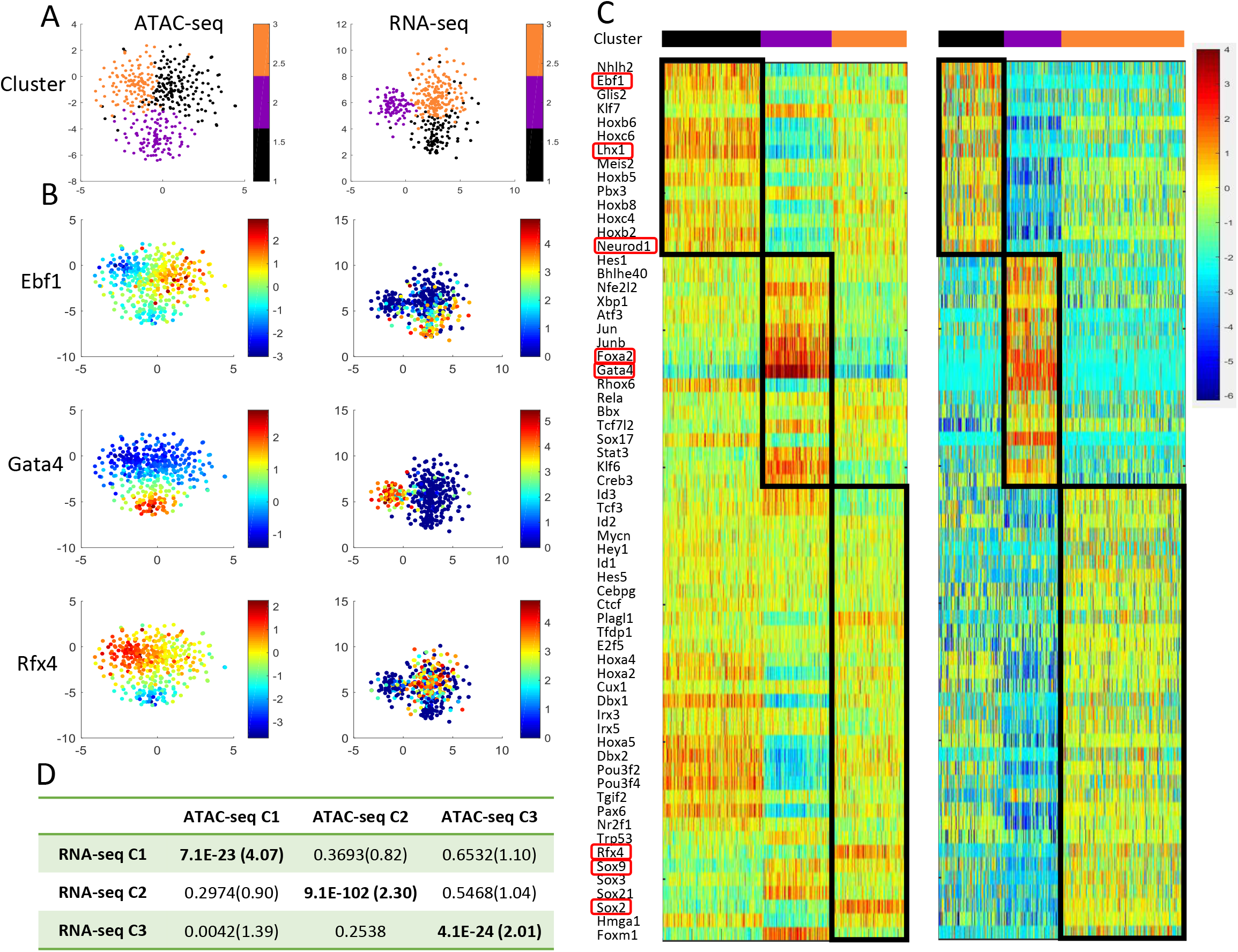
(A) t-SNE plot of single cell RNA-seq (right) and single cell ATAC-seq data (left) from RA day 4. Different colors represent clustering assignment from coupled clustering method. (B) Same t-SNE plots as Fig. 3 A. Different colors represent cluster-specific TFs’ (Ebf1, Gata4, and Rfx4) gene expression Z-score and motif activity Z-score. (C) Comparison of cluster-specific TFs’ expression Z-score with motif activity Z-score on cluster level. (D) Overlap of cluster-specific peaks nearby genes with cluster-specific genes. The values in table represent fisher’s exact test p-value and fold change.

Next, we select cluster-specific genes from RNA-seq data and cluster-specific peaks from ATAC-seq data. We assess whether the cluster-specific peaks are significantly close to the cluster-specific-genes by performing Fisher exact test based on the count of such gene-peak pairs that are within 100kb of distance each other (Supplementary Fig. S5). Figure 4D gives the p-values for all possible pairings of the RNA-seq clusters with ATAC-seq clusters. It is seen that the pairings identified by coupled NMF indeed gave dramatically more significant p-values and higher fold changes than the other possible pairings.

### Coupled clustering of single cells sheds light on stem cell differentiation

The cluster-specific gene expression profiles and chromatin accessibility profiles provided by our method can provide useful insight on the constituent subpopulations. First, we use cluster-specific peaks from single cell ATAC-seq data to annotate the clusters. We collect previously determined enhancers in mouse tissues at 7 developmental stages from 11.5 days post conception until birth (10). Fig. 5 *A-C* shows the degree of overlap of our cluster-specific peaks with these developmental enhancers for different tissues and at different developmental stages. The number represents 10,000 times Jaccard index (intersection over union) and NA indicates that enhancer data for that tissue in that stage is not available. The results show that cluster-1-specific peaks are enriched in forebrain and midbrain enhancers at E12.5 and E13.5. Cluster-2-specific peaks are enriched in heart enhancers at E15.5 and E16.5. Cluster-3-specific peaks are enriched in forebrain enhancers from E12.5 to E16.5 and also in midbrain, hindbrain and neural tube. We also collect experimentally validated tissue-specific enhancers from VISITA database and overlap them to cluster-specific peaks. Fig. 5 *D*shows the percentage of tissue specific VISITA enhancers overlapped to cluster-specific-peaks. Only those tissues with at least one enhancers overlapping with the cluster-specific peaks are shown. Enhancers from neural associated tissue (neural tube, cranial nerve, hind brain, midbrain, forebrain, trigeminal V, dorsal root ganglion, eye, nose) have overlap with cluster-specific peaks from cluster 1 and cluster 3. Cluster 2 specific-peaks are enriched in blood vessels enhancers and heart enhancers. These results suggest that cluster 1 and 3 may be related to neural tissues and cluster 2 may be related to heart tissue.

**Fig. 5.**
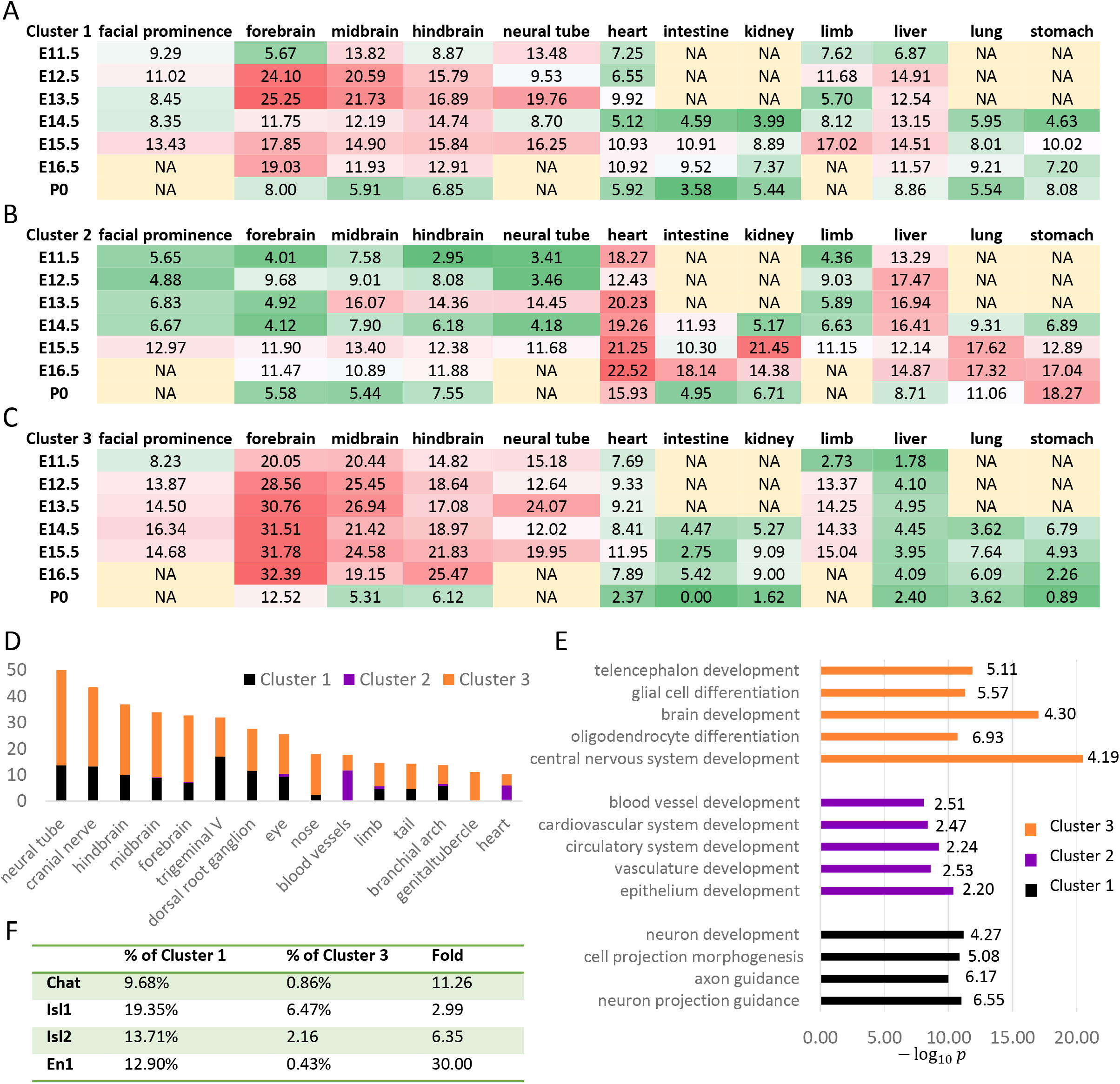
(A-C) Similarity of cluster-specific peaks with enhancers of 12 tissues’ 7 developmental stages. The number represent 10,000 times Jaccard index and NA represent enhancer data of that tissue in that stage are not available. (D) Percentage of VISITA enhancer that overlapped with cluster-specific peaks. (E) GO enrichment of cluster-specific genes. (F) Comparison of motor neuron markers’ expression on cluster 1 and 3. Number represent the percentage of expressed cells (TPM >2) on cluster.

In addition, we analyzed cluster-specific genes from scRNA-seq data. Fig. 5 *E* presents the most enriched gene ontology (GO) terms, their p-values and fold changes in each cluster. The results show that cluster 2 is enriched in blood vessel development and cardiovascular system development, cluster 1 and 3 are enriched in neuron associated terms. The results from scRNA-seq based annotation are consistent with the results from scATAC-seq based analysis. Although clusters 1 and 3 are neural associated clusters, there are interesting differences. Cluster 1 is more enriched in axon guidance and neuron projection guidance, which are relevant for general neuronal functions. On the other hand, cluster 3 are more enriched in brain development and oligodendrocytes differentiation, which are specifically relevant to the central nervous system (CNS). To examine this further, we check the expression of some known markers for motor neurons, which are present in PNS (peripheral nervous system) rather than in CNS. Fig. 5F shows the percentage of cells expressing (i.e. TPM>2) motor neuron markers (Chat, Isl1, Isl2 and En1). Motor neuron markers are more highly expressed in cluster 1 compared to cluster 3. Overall our results suggest that retinoic acid induced stem cell at day 4 is a mixture of cells related to peripheral nervous system, cardiovascular system, and central nervous system. These results are largely consistent with the previous studies (11, 12).

We can construct cluster-specific gene regulatory networks as graphs with directed edges from the cluster-specific peaks to the cluster-specific genes that are within 100 kb distance, and directed edges from cluster-specific TFs to cluster-specific peaks containing significant matches to the corresponding motifs. These cluster-specific subnetworks are presented in Supplementary Fig. S6. It is seen that Klf7, Ebf1, Sox11, and Nhlh1 are playing important role in the network for cluster 1, Gata4, Gata6, Sox17, Foxa2, Ap1 complex and Tead family are important in cluster 2 and Rarb, Nr2f1, Rfx4, Sox2, Sox9, Sox21, Pou3f2, Pou3f3, and Pou3f4 are important in cluster 3.

## Discussions

As far as we know, coupled clustering is a new problem different from other complex clustering tasks such as bi-clustering or multi-view clustering. Bi-clustering (13, 14), also called block clustering or co-clustering, has been used widely used to cluster subjects and cluster genes simultaneously based on a *p* by *n* data matrix of expression measurements on *p* genes for *n* subjects. The same data matrix is used the clustering in gene space as well as the clustering in subject space. In contrast, two different data matrices are used in coupled clustering of two separate samples. In multi-view clustering (15), the set of features measured on each subject can be divided into two independent subsets, for example, one of them may represent gene expression measurements while the other represent accessibility measurements. The important difference between multi-view clustering and coupled clustering is that in the former setting all features are measured on each subject, whereas in the latter only one of the subsets can be measured on any subject. Clearly, coupled clustering is a more challenging task and requires external information such as subject domain knowledge or prior data in order to initialize the coupling.

Measurements of different types of features generally require different types of reagents and biochemical reactions. To measure multiple types of features in the same cell, it is necessary to carry out all the required reactions within the same cell, which is extremely challenging technically. Although there are current efforts to develop methods for simultaneous measurement of two types of measurement in single cells (for example, RNA+DNA, RNA+accessibility, or RNA+CpG methylation, etc), these methods are still not yet ready for general use. Furthermore, even when we have two types of data (say RNA + accessibility) on single cells, there are always additional features (say CpG methylation) that we may want to incorporate into the analysis. For example, we may have two types of features measured in single cells in the first sample, and a third type of features measured on single cells in another sample. Then we are again facing a coupled clustering problem. Thus our method for coupled clustering is of interest to single cell genomics.

## Methods and Materials

### Optimization algorithm

We optimize the object function in *eq. (3) by* multiplicative update algorithm.

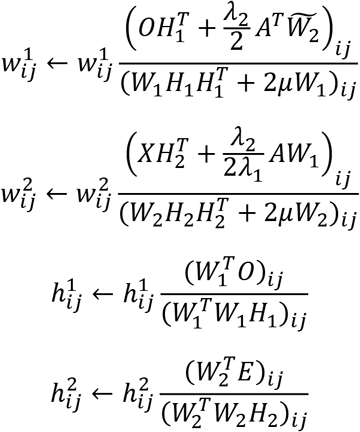

Where 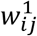 represent the element of *i^th^* row and *j^th^* column in matrix *W*_1_, The same representation is used in *W*_2_, *H*_1_ and *H*_2_. We stop the iteration when the relative error is less than 0.0001.

### Cluster-specific features

We apply t-test to define the cluster-specific genes and cluster-specific peaks, default p-value cut-off is 0.0001.

### Evaluation of the clustering results

We evaluate the results in terms of consistency of true expression values and the predicted values. We calculated the correlation *K* × *K* matrix of *AW_1_* with 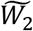, which is denoted by *R*. We use the determinant of correlation matrix *R* to measure the consistency of true expression values and the predicted values. Higher determinant means higher diagonal of the matrix, which means higher correlation between matched clusters and lower correlation in unmatched clusters.

### Parameters selection

We solve optimization problems 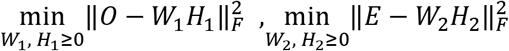 by alternating least squares (ALS) algorithm with 50 different initializations using a Monte Carlo type approach (16) and get the solutions *W*_10_, *H*_10_, *W*_20_, *H*_20_, which are used as initial solution in our optimization problem. We choose parameters 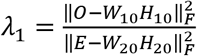, 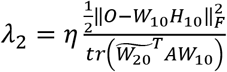 Tuning parameters *η* and *μ* are chosen from 0.001, 0.01, 0.1, 1, 10, 100, 1,000, and 10,000. The determinant of correlation matrix *R* can be used to select the tuning parameters. We choose the tuning parameters which give the highest determinant. The number of clusters *K* can be determined by a method similar to that in ref (17).

### TF motif activity

We use software chromVAR (18) to calculate the TF motif activity on each of the scATAC-seq data.

### Single cell sample at RA day-4

We generated a heterogeneous biological population of cells that arise from the same origin. Specifically, we used the hanging drop technique to form embryonic bodies (EBs) from mouse embryonic stem cells (mESCs) and induced differentiation by retinoic acid (RA) treatment. After 4 days’ induction, we sample cells for bulk RNA-seq and bulk ATACC-seq experiments for use in validating the coupling. To test the couple NMF clustering method, we also generated single cell ATAC-seq and single cell RNA-seq on the RA day-4 population. After removing low read count cells (3,000 in RNA-seq and 10,000 in ATAC-seq), we get ATAC-seq data and RNA-seq data on 415 and 463 single cells respectively.

### Data processing

We align the single cell ATAC-seq reads to reference genome mm9 and remove duplicates. We employed MACS2 (19) to do peak calling by merging all the reads from all the single cells. We only consider the narrow peaks which at least present (1 or more reads) on 10 cells. Read counts for each region on each cell are calculated by bedtools (20) with intersect command. Feature of scATAC-seq data is regions including regulatory elements (REs) and narrow peaks from MACS2. REs include promoters and enhancers. We use REs that regulate at least one target gene from PECA network (8).

Single cell RNA-seq raw reads are mapped to mm10 by STAR (21) with ENCODE options. Gene expression transcripts per million (TPM) are calculated by RSEM (22). The transcriptome annotation we use is GENCODE vM16.

### Experimental design of retinoic acid-induced mESC differentiation

Mouse ES cell lines R1 were obtained from ATCC. The mESCs were first expanded on an MEF feeder layer previously irradiated. Then, subculturing was carried out on 0.1% bovine gelatin-coated tissue culture plates. Cells were propagated in mESC medium consisting of Knockout DMEM supplemented with 15% Knockout Serum Replacement, 100 μM nonessential amino acids, 0.5 mM beta-mercaptoethanol, 2 mM GlutaMax, and 100 U/mL Penicillin-Streptomycin with the addition of 1,000 U/mL of LIF (ESGRO, Millipore).

mESCs were differentiated using the hanging drop method (23). Trypsinized cells were suspended in differentiation medium (mESC medium without LIF) to a concentration of 50,000 cells/ml. 20 μl drops (~1000 cells) were then placed on the lid of a bacterial plate and the lid was upside down. After 48 h incubation, Embryoid bodies (EBs) formed at the bottom of the drops were collected and placed in the well of a 6-well ultra-low attachment plate with fresh differentiation medium containing 0.5 μM retinoic acid (RA) for up to 4 days, with the medium being changed daily.

### Single cell ATAC-seq

We followed the single cell ATAC-seq protocol published by Jason et al. (24) with the following modifications. The EBs were first incubated with StemPro Accutase cell dissociation reagent (Gibco) at 37°C for 10 min, then the EBs were gently pipetted for additional 15 min to obtain single cell suspension. To further remove non-dissociated EBs, the cell suspension was filtered sequentially with a 40 μM cell strainer (BD Falcon) and a 20 μM pluriStrainer (pluriSelect). After washing 3 times with C1 DNA Seq Cell Wash Buffer, cells at a concentration of 350-400 cells/μl were loaded on the C1 Single-Cell Auto Prep System (Fluidigm, Inc.). Single cells were captured and processed on a 10-17 μM IFC microfluidic chip using ATAC-seq scripts (24). Total 7 IFC chips were included in this study. The library was sequenced on Illumina NextSeq with 75 bp paired-end reads.

### Single cell RNA-seq

To prepare single cell RNA-seq library, we followed SMART-Seq v4 Ultra Low Input RNA Kit for the Fluidigm C1 System (Clontech Laboratories, Inc.). The EBs were first dissociated with Accutase as described previously. Cells at a concentration of 200-250 cells/μl were then loaded on the C1 Single-Cell Auto Prep System (Fluidigm, Inc.). The single cells were captured and processed on a 10-17 μM IFC microfluidic chip using SMART-Seq v4 scripts. Total 5 IFC chips were included in this study. After harvest, cDNA concentration for each sample was measured using the Fragment Analyzer Automated CE System (Advanced Analytical Technologies, Inc.) and the cDNA concentration we used for Nextera XT library preparation is ~0.2 ng/μl. The library was sequenced on Illumina HiSeq with 100 bp paired-end reads.

### Software and Data

Software and simulation data are available at web.stanford.edu/~zduren/CoupledClustering/. Single cell gene expression data and chromatin accessibility data of RA induction have been deposited in the GEO database under accession no. GSEXXXXX.

## Acknowledgments

This work was supported by grants R01HG007834, R01GM109836, and P50HG007735 from the National Institutes of Health (NIH).

## Supporting information

**Supplementary Figure S1.**
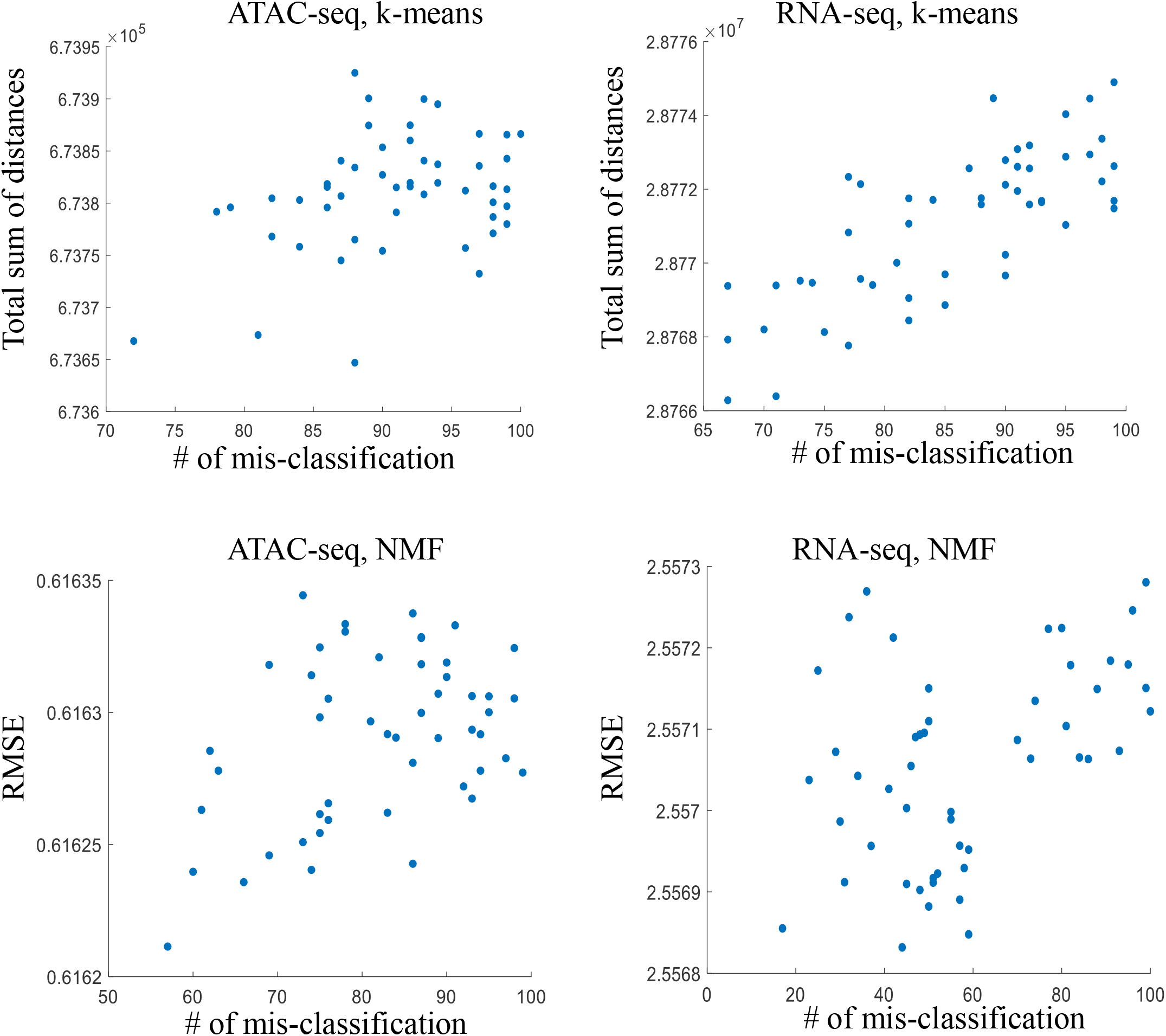
Clustering results of k-means and NMF on simulation data. Each dot represents a run with random initial.

**Supplementary Figure S2.**
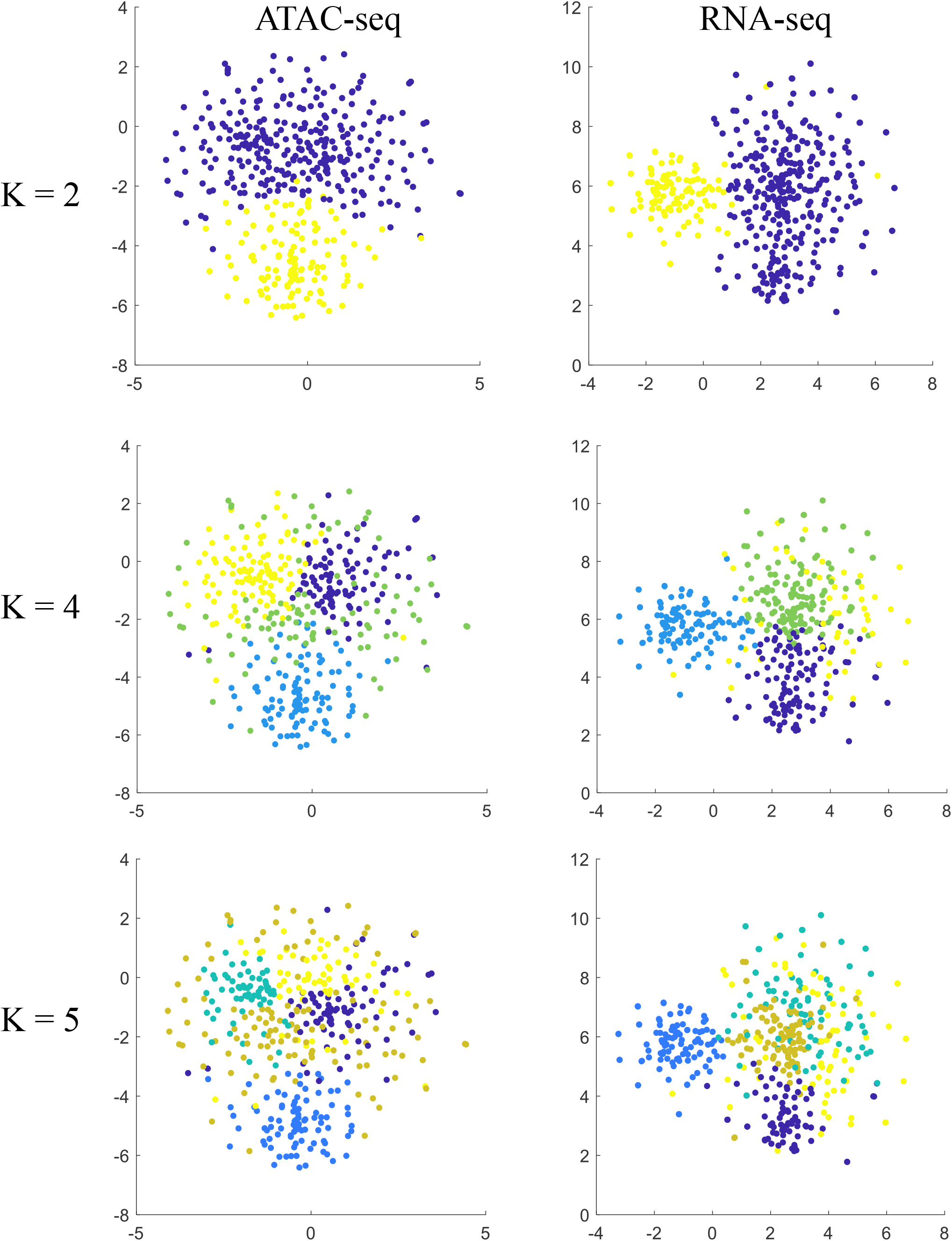
t-SNE plot of scATAC-seq and scRNA-seq data on RA day 4 sample. Each dot represents one cell and different colors represent different clusters.

**Supplementary Figure S3.**
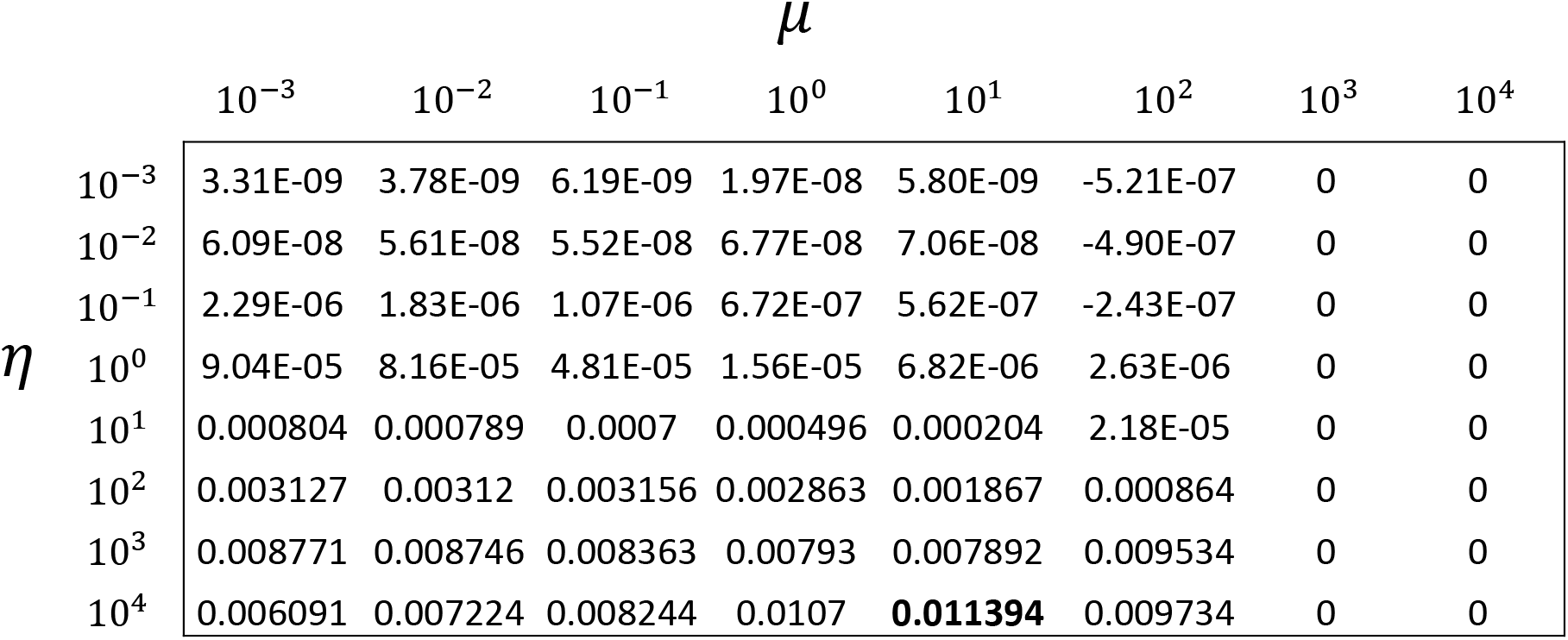
Selecting tuning parameters *μ* and *η*. The values represent the determinant of correlation matrix R=corr (AW_1_,W_2_).

**Supplementary Figure S4.**
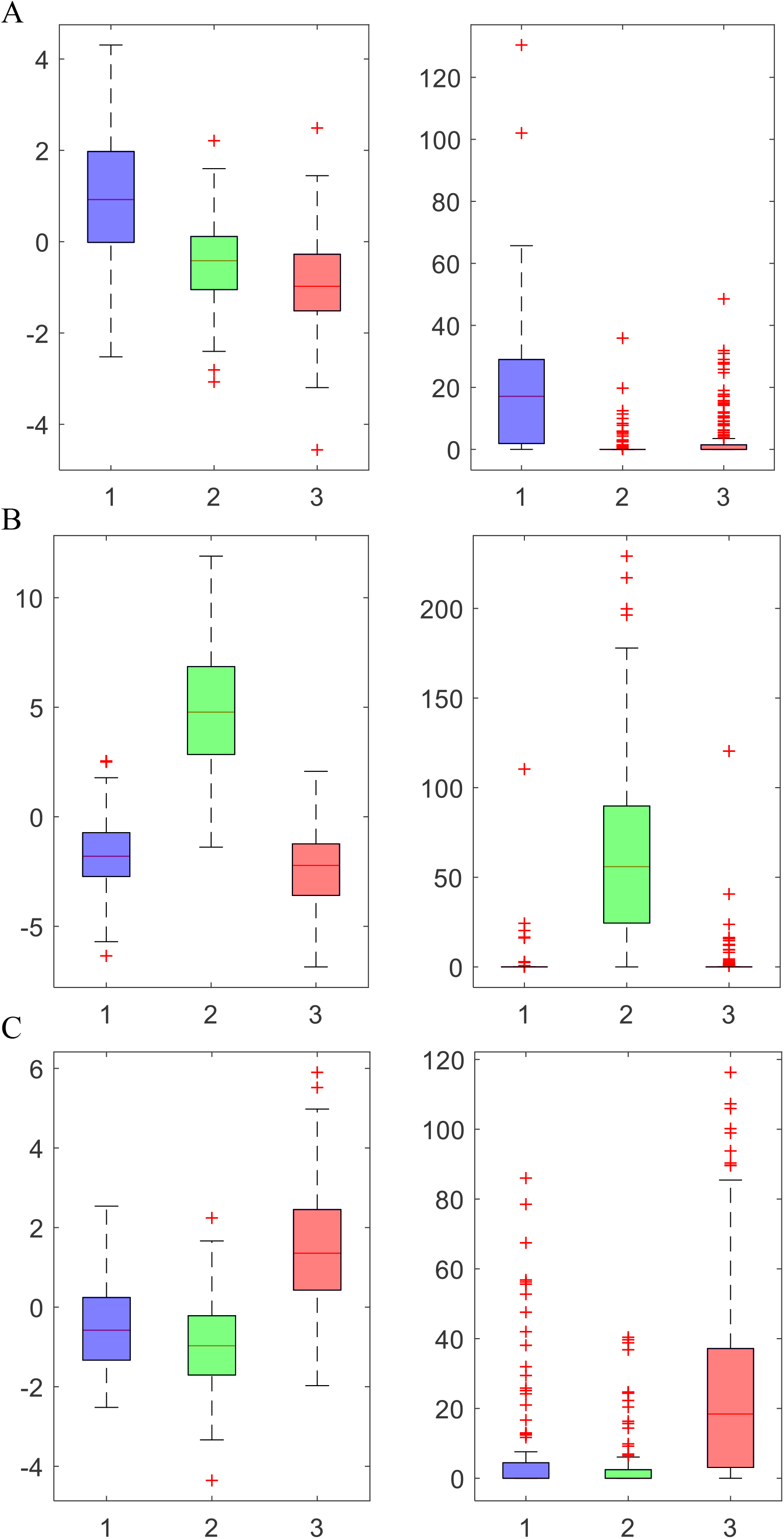
Boxplot of cluster specific markers’, Ebf1, Gata4, and Rfx4, motif activity (left) and gene expression (right).

**Supplementary Figure S5.**
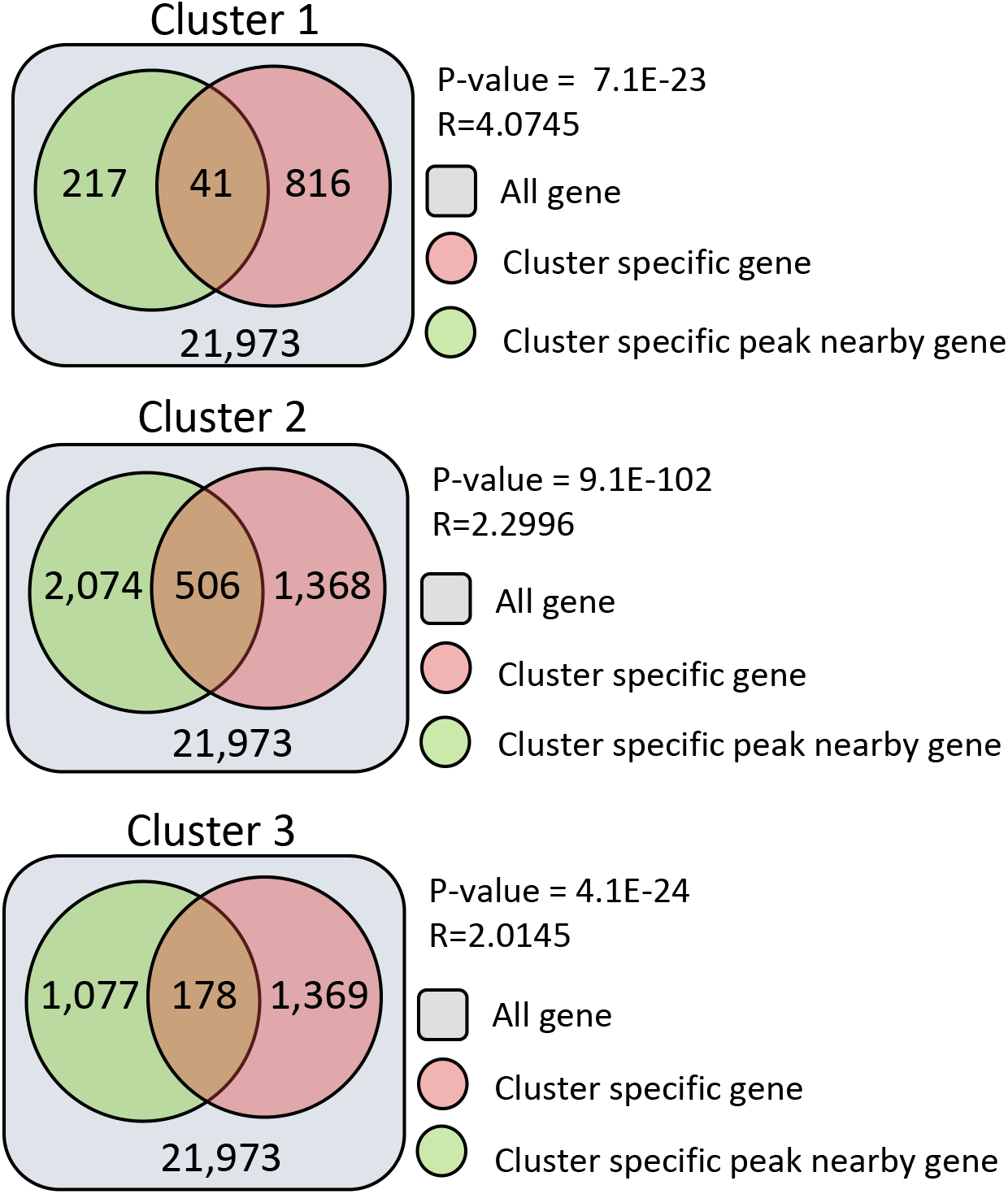
Comparison of cluster specific genes with the cluster specific peak nearby gene on each cluster from RA day 4 data.

**Supplementary Figure S6.**
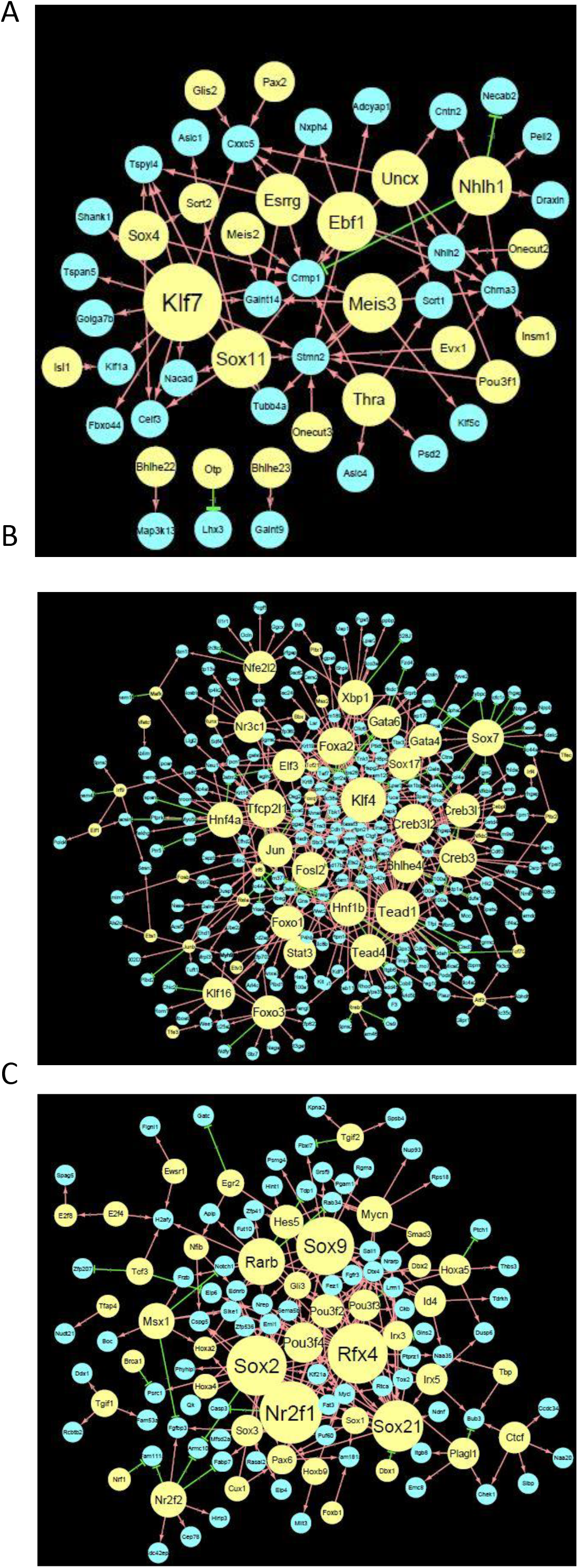
(A-C) Cluster specific networks of cluster 1, 2, and 3. Yellow color nodes represent TFs, red edges represent activation, and green edges represent repression. Circle size is proportional to the out-degree.

## Supplementary Table legends

Supplementary Table 1. List of human and mouse paired gene expression and chromatin accessibility data used in regression model.

